# MicroRNA-1 promotes the development of and prolongs engorgement time in *Hyalomma anatolicum anatolicum* (Acari: Ixodidae) ticks

**DOI:** 10.1101/2020.08.19.257014

**Authors:** Jin Luo, Qiaoyun Ren, Wenge Liu, Xiaofei Qiu, Gaofeng Zhang, Yangchun Tan, Runlai Cao, Hong Yin, Jianxun Luo, Xiangrui Li, Guangyuan Liu

## Abstract

MicroRNAs act as mRNA posttranscriptional regulators, playing important roles in cell differentiation, transcriptional regulation, growth and development. In this study, microRNA expression profiles of *Hyalomma anatolicum anatolicum* ticks at different developmental stages were detected by high-throughput sequencing and functionally assessed. In total, 2,585,169, 1,252,678, 1,558,217 and 1,155,283 unique reads were obtained from eggs, larvae, nymphs and adults, respectively, with 42, 46, 45 and 41 conserved microRNAs in these stages, respectively. Using eggs as a control, 48, 43 and 39 microRNAs were upregulated and 3, 10 and 9 downregulated in larvae, nymphs and adults, respectively. microRNA-1 (miR-1) was expressed in high abundance throughout *Ha. anatolicum* development, with an average of nearly one million transcripts, and it is highly conserved among tick species. Quantitative real-time PCR (qPCR) showed that miR-1 expression gradually increased with tick development, reaching the highest level at engorgement. Differential tissue expression was detected, with significantly higher levels in the salivary glands and epidermis than in the midgut. Inhibition assays showed no significant change in body weight or spawning time or amount between experimental and control groups, but there was a significant difference (p<0.01) in engorgement time. With miR-1 inhibition, ticks displayed obvious deformities during later development. To more fully explain the microRNA mechanism of action, the miR-1 family was analyzed regarding target gene; members acting on Hsp60 include miR-5, miR-994, miR-969, and miR-1011, which jointly play a role. Therefore, microRNAs are critical for normal tick development, and the primary structure of the mature sequence of miR-1 is highly conserved. Nonetheless, different developmental stages and tissues show different expression patterns, with a certain role in prolonging feeding. miR-1, together with other family members, regulates mRNA function and may be used as a molecular marker for species origin and evolution analysis and internal reference gene selection.

## Introduction

*Hyalomma anatolicum anatolicum* belongs to taxa Parasitiformes, Ixodoidea, Ixodidae and Hyalomma. At present, ticks are only reported in semidesert areas of Xinjiang Uyghur autonomous region in China [1]. However, ticks are distributed worldwide, including the former Soviet Union, India, Nepal, Pakistan and Central Asia, North Africa and eastern European countries. Ticks parasitize cattle, sheep, camels, horses, donkeys and a few wild animals, serve as a transmission vector of *Crimean-Congo hemorrhagic fever* (Xinjiang hemorrhagic fever), can be naturally infected with *Coxiella burnetii* (Q fever), and can spread brucellosis and piriformis disease [2,3]. Ticks cause serious harm to the livestock industry and public health.

As key components of most regulatory events, microRNAs play important roles at the posttranscriptional level in various developmental and physiological processes [4–12]. MicroRNA-1 (miR-1) is another important microRNA molecule discovered in recent years. Its precursor molecules regulate target genes in cells. Studies have confirmed two subtypes of mature miR-1 (miR-1-1 and miR-1-2) in primates. These microRNAs play an important role in skeletal and cardiac muscle development [13,14]. In particular, miR-1 is crucial in the development of cardiac hypertrophy, myocardial infarction, arrhythmia and other cardiac diseases [15–17]. At present, downregulation of miR-1 has been widely used as a biomolecular marker for myocardial infarction [18–23]. Other studies have shown that heat shock protein 60 (Hsp60) is a target gene of miR-1.

Hsp60 is a component of the defense mechanism against diabetic myocardial injury. The expression level of Hsp60 is significantly decreased in diabetic myocardial injury tissues. In vivo and in vitro experiments confirm that increased glucose levels in atrophic cardiomyocyte cells lead to upregulation of miR-1 expression, which accelerates glucose-mediated apoptosis by Hsp60 in cardiomyocytes [24]. miR-1 also plays an important role in the differentiation and development of smooth muscle and skeletal muscle cells [25–27]. For example, miR-1 was found to be a factor in specifically differentiated smooth muscle cells isolated from embryonic stem cell-derived cultures. Loss of miR-1 function can cause a decrease in smooth muscle biomarkers and the number of derived smooth muscle cells. Indeed, evidence has shown that miR-1 is the main factor regulating smooth muscle cell differentiation, and Kruppel-like 4 (KLF4) is affected by miR-1 downregulation. The recognition site of miR-1 is located in the 3’-UTR of KLF4, and inhibition of miR-1 reduces KLF4 expression and smooth muscle cell differentiation [28].

Mutation of mir-1 and mir-206 sites in the 3’-UTR of muscle growth inhibition factor have been established in the Texel sheep model, leading to changes in muscle phenotype [29]. Therefore, it is important to understand miR-1 function by examining targets of microRNA families. There are 142 target genes of the miR-1 family based on Pictar software and 187 conserved sites and 14 nonconservative sites with 181 conserved genes according to TargetScanFly. Gene Ontology enrichment analysis shows that target genes of the miR-1 family are involved in various biological processes, including gene expression regulation, nucleic acid metabolism, cell communication, cell division, growth and proliferation. Therefore, studying the function of target genes regulated by miR-1 in genetic interaction networks is of great significance for better understanding the mechanisms of certain diseases and drug development.

miR-1 plays an important regulatory role in mammalian muscles, but its function in ticks remains unclear. Here, the expression level of miR-1 in different developmental stages and tissues of ticks is analyzed to predict its possible biological functions. An inhibitor of miR-1 was injected into the fourth sarcomere, and physiological indicators were assessed at various developmental stages. The molecular mechanisms were also investigated, and the biological function of miR-1 in tick development was explored. This is the first report on the function of miR-1 in ticks, and we present the characteristics of tick development upon abnormal expression of miR-1. This study provides a new insight into the function of microRNAs in ticks as well as a theoretical basis for the prevention and control of ticks.

## Materials and Methods

### Ethics statement

No ethics approvals are required for work with *Ha. anatolicum*.

### Tick collection and RNA extraction

In this study, *Ha. anatolicum* ticks were obtained from Xinjiang Uyghur Autonomous Region (XUAR) and identified using morphology by the Animal Research Institute (Lanzhou Veterinary Research Institute). After cultivation, eggs were collected into sterile tubes for incubation. Approximately 2 g of larvae, on average, were divided into two parts. One batch was obtained for microRNA extraction. The other batch was placed in a mesh bag attached to a host for 27 days until the unfed nymphs and unfed adult ticks were collected.

To analyze the dynamic characteristics of microRNAs from different developmental stages of *Ha*. *anatolicum,* four samples (eggs, larvae, nymphs and adults) were homogenized by freezing in liquid nitrogen and grinding into a powder using a sterile mortar and pestle. Total RNA and/or enriched small RNA fractions were isolated from whole-tick samples using the miScript microRNA isolation kit according to the manufacturer’s instructions (QIAGEN, China). The quantity and integrity of the total RNA was assessed using an Agilent 2100 Bioanalyzer system (Agilent Technologies, USA). Total RNA was stored at −80°C until use.

### Small RNA isolation and high-throughput sequencing

The quality of total RNA was analyzed using a Shimadzu 206-97213C BioSpec-nano analyzer system with denaturing polyacrylamide gel electrophoresis. A small RNA library was generated according to the Illumina sample preparation instructions [30]. Briefly, total RNA samples were size-fractionated on a 15% Tris-borate-EDTA-urea polyacrylamide gel. RNA fragments 18–50 nt long were isolated, quantified, and ethanol-precipitated. A 5’-adapter (Illumina) was ligated to the RNA fragments with T4 RNA ligase (Promega). The ligated RNAs were size-fractionated on a 15% Tris-borate-EDTA-urea polyacrylamide gel, and 41 ~ 76-nt-long RNA fragments were isolated. Next, 3’-adapter (Illumina) ligation was performed, followed by a second size fractionation using the same gel conditions as described above. The 64 ~ 99-nt-long RNA fragments were isolated by gel elution and ethanol precipitation. The ligated RNA fragments were reverse-transcribed to single-stranded cDNAs using M-MuLV (Invitrogen) with RT primers (as recommended by Illumina). cDNAs were amplified with pfx DNA polymerase (Invitrogen) using 20 PCR cycles and the Illumina small RNA primer set. The PCR products were purified on a 12% Tris-borate-EDTA polyacrylamide gel, and a slice containing cDNAs of 80~115 bp was excised. This fraction was eluted, and the recovered cDNAs were precipitated and quantified using a Nanodrop (Thermo Scientific) and TBS-380 minifluorometer (Turner Biosystems) with PicoGreenH dsDNA quantization reagent (Invitrogen). The concentration of the sample was adjusted to 10 nM, and 10 μL was used for sequencing. The purified cDNA library was used for cluster generation (with the Illumina Cluster Station) and then sequenced using a HiSeq2000 following the manufacturer’s instructions.

### Small RNA bioinformatics analysis

Sequence data (raw data or raw reads) conversion was conducted by base calling. We used software developed by BGI for HiSeq sequencing data processing, eliminating some contaminants and low-quality reads to obtain final clean reads. The data were processed according to the following steps: 1) removing low-quality reads; 2) removing reads with 5’ primer contamination; 3) removing reads without a 3’ primer; 4) removing reads without the insert tag; 5) removing reads with poly-A; 6) removing reads shorter than 18 nt; 7) summarizing the length distribution of the clean reads. Normally, the length of a small RNA is between 18 nt and 30 nt, and length distribution analysis is helpful to assess the length compositions of a small RNA sample. For example, microRNA is normally 21 nt or 22 nt, siRNA is 24 nt, and piRNA is 30 nt. The clean read data were assembled using SOAPdenovo short sequence assembly software (http://soap.genomics.org.cn/soapdenovo.html:1.05) and used to assemble small RNAs for mapping to the *Ixodes scapularis* genome by bowtie software (http://bowtie-bio.sourceforge.net/manual.shtml) and SOAP assembly [31]. Small RNAs were aligned to the microRNA precursor of corresponding species (the mature microRNA if no precursor information for that species was found in miRBase21) to obtain the microRNA count as well as base bias at the first position of the identified microRNAs with certain lengths and for each position of all identified microRNAs. The small RNA tags were annotated as rRNA, scRNA, snoRNA, snRNA or tRNA using GenBank and Rfam databases with tag2 annotation software (developed by BGI). In the above alignment and annotation, some small RNA tags may be mapped to more than one category. Thus, to ensure unique small RNAs mapped to only one annotation, we followed the following priority rule: rRNAetc (in which GenBank > Rfam) > known microRNA > repeat > exon > intron [32]. The total rRNA proportion is a marker of sample quality control, whereby high-quality samples should be less than 60% for plants and 40% for animals. The unannotated sequences were used to predict potential novel microRNA candidates.

The characteristic hairpin structure of the microRNA precursor can be used to predict novel microRNAs [33]. BGI developed the prediction software Mireap to predict novel microRNA by exploring the secondary structure, Dicer cleavage site and minimum free energy of the unannotated small RNA tags mapped to a genome. Mireap can be accessed at http://sourceforge.net/projects/mireap/ [34] under the following parameter settings according to Zuker and Jacobson [35]: minimal microRNA sequence length 18; maximal microRNA sequence length 26; minimal microRNA reference sequence length 20; maximal microRNA reference sequence length 24; minimal depth of Drosha/Dicer cutting site 3; maximal copy number of microRNAs on reference 20; maximal free energy allowed for a microRNA precursor −18 kcal/mol; maximal space between microRNA and microRNA* 35; minimal base pairs of microRNA and microRNA* 14; maximal bulge of microRNA and microRNA* 4; maximal asymmetry of microRNA/microRNA* duplex 5; flanking sequence length of microRNA precursor 10. Stem-loop hairpins were considered typically in accordance with the following three criteria [35]: mature microRNAs were present in one arm of the hairpin precursors, which lack large internal loops or bulges; the secondary structures of the hairpins were stable, with a free energy of hybridization lower than −18 kcal/mol; and hairpins were located in intergenic regions or introns.

Genes with sequences and structures that fulfilled the three criteria, forming perfect stem-loop structures, were considered microRNA candidates. Finally, all remaining novel microRNA candidates were subjected to MiPred (http://www.bioinf.seu.edu.cn/microRNA/) to filter out pseudopremicroRNAs using the following settings: minimum free energy >−20 kcal/mol or P-value>0.05 [36].

### Prediction of microRNA targets and GO analysis

Because no 3’-UTR database is currently available, putative target genes of novel microRNA candidates were explored by aligning microRNA sequences with the tick EST database in NCBI. The rules used for target prediction were based on those suggested by Allen (Allen E, 2005) and Schwab (Schwab R, 2005), as follows: (1) no more than four mismatches between sRNA and target (G-U bases count as 0.5 mismatches); (2) no more than two adjacent mismatches in the microRNA/target duplex; (3) no adjacent mismatches in positions 2~12 of the microRNA/target duplex (5’ of microRNA); (4) no mismatches in positions 10~11 of microRNA/target duplex; (5) no more than 2.5 mismatches in positions 1 ~ 12 of the of the microRNA/target duplex (5’ of microRNA); and (6) minimum free energy (MFE) of the microRNA/target duplex be ≥ 75% of the MFE of the microRNA bound to its perfect complement. More strictly, no more than two mismatches between the microRNA sequence and potential microRNA target were allowed.

Gene Ontology (GO) is an international standardized classification system for gene function that supplies a set of controlled vocabulary to comprehensively describe the properties of genes and gene products. There are 3 ontologies in GO: molecular function, cellular component and biological process. GO terms significantly enriched for the predicted target gene candidates of microRNAs compared with the reference gene background and the genes corresponding to certain biological functions were analyzed. This method first maps all target gene candidates to GO terms in the database (http://www.geneontology.org/) [37], calculating gene numbers for each term, and then applies a hypergeometric test to find significantly enriched GO terms for target gene candidates compared to the reference gene background.

### Real-time quantitative PCR

Stem-loop real-time reverse transcription polymerase chain reaction (RT-PCR) with SYBR Green was used for the analysis of microRNA expression in *Ha*. *anatolicum* according to the manufacturer’s protocol. A stem-loop forward primer (5’-GTC GTA TCC AGT GCA GGG TCC GAG GTA TTC GCA CTG GAT ACG AC-3’) was used to quantify microRNA expression because it can provide more specificity and sensitivity than linear primers. The reverse universal primer (10×miScript Universal primer) was provided by QIAGEN Co, Ltd, China, and the forward primer was designed by Primer Premier 5.0 (miR1F: 5′-TCC GTT CGG ATC ACC GTG CTT C-3′). The *β*-*Actin* (EF488512) gene was designed as a reference gene, with the following primers used: sense primer, 5′-TGT GAC GAC GAG GTT GCC G-3′; anti-sense primer, 5′-GAA GCA CTT GAG GTG GAC AAT G-3′. All forward primers were synthesized by Shenggong Co, Ltd, China (Table 1). Real-time quantitative PCR was performed using Mx3000pTM SYBR Green real-time quantitative PCR Analyzer (QIAGEN Biotechnology Co, Ltd, China). Briefly, 2 μg of microRNA was reverse transcribed using miScript Ⅱ microRNA cDNA Synthesis Kit (QIAGEN Biotechnology Co, Ltd, China). The reverse transcription reaction system included 4 μL of 5×miScript HiFlex Buffer, 2 μL of 10×miScript Nucleics Mix, 2 μL of miScript Reverse Transcriptase Mix and RNase-free dH_2_O to a final volume of 20 μL. The RT-PCR program was set to 37°C for 60 min followed by 95°C for 5 min. The cDNA products were stored at −20°C. Relative real-time quantitative PCR was performed with a miScript SYBR Green PCR kit (QIAGEN Biotechnology Co, Ltd, China). The reaction solution was prepared on ice and comprised 10 μL of 2× QuantiTect SYBR Green PCR Master Mix, 2 μL of 2× miScript Universal primer (10 μM), 2 μL of forward primer (10 μM), 2 μL of cDNA, and dH_2_O to a final volume of 20 μL. The reaction mixtures were incubated in a 96-well plate at 95°C for 15 min, followed by 35 cycles of 94°C for 15 sec, 60°C for 30 sec and 70°C for 30 sec. All reactions were performed in triplicate. The primers for the microRNAs had the same sequences as the tick microRNAs with appropriate adjustments at their 5′ terminus. Mx3000/Mx Pro software (Stratagene, USA) was used to construct a melting curve. Standard curves with 5-fold dilutions were performed for each assay, and PCR efficiency calculations were based on the slopes of the standard curves. The absolute amount of each microRNA was calculated using the 2^−△△CT^ method [38] according to the standard curve. The housekeeping gene U6 was employed as an endogenous control, and the U6 primers were provided by QIAGEN Co, Ltd, China. Each sample was replicated three times. The microRNA level in various samples of developmental stages was determined individually. Each microRNA level is expressed as the 2^−△△CT^ mean ± SE. One-way ANOVA was applied to examine the significance of differential expression level in each mature/novel microRNA between eggs and larvae, larvae and adults, eggs and adults, and the difference was considered significant at P<0.05. Clones containing an insert of the correct size from four independent PCRs were sequenced on both strands using an ALF sequencer (Pharmacia Biotech).

**Table 1.** Alignment to Genbank and Rfam data librarys. Annotate the sRNA tags with rRNA, scRNA, snoRNA, snRNA and tRNA from 13 Genbank by BLAST. For some species, its non-coding RNAs from Genbank need to be supplied as reference for this analysis.

| Category | Egg |  |  |  | Larvae |  |  |  | Nymph |  |  |  | Adult |  |  |  |
| --- | --- | --- | --- | --- | --- | --- | --- | --- | --- | --- | --- | --- | --- | --- | --- | --- |
|  | Unique | Percent(%) | Total | Percent(%) | Unique | Percent(%) | Total | Percent(%) | Unique | Percent(%) | Total | Percent(%) | Unique | Percent(%) | Total | Percent(%) |
|  | sRNAs |  | sRNAs |  | sRNAs |  | sRNAs |  | sRNAs |  | sRNAs |  | sRNAs |  | sRNAs |  |
| Total | 2,585,169 | 100 | 16,262,023 | 100 | 1,252,678 | 100 | 9,835,555 | 100 | 1,558,217 | 100 | 9,202,455 | 100 | 1,155,283 | 100 | 9,147,849 | 100 |
| miRNA | 33,311 | 1.29 | 987,808 | 6.07 | 35,866 | 2.86 | 1,483,548 | 15.08 | 26,866 | 1.72 | 799,098 | 8.68 | 22,193 | 1.92 | 1,627,034 | 17.79 |
| rRNA | 41,612 | 1.61 | 1,322,480 | 8.13 | 58,098 | 4.64 | 602,546 | 6.13 | 33,881 | 2.17 | 772,142 | 8.39 | 23,876 | 2.07 | 1,023,248 | 11.19 |
| repeat | 6 | 0.00 | 6 | 0.00 | 41 | 0.00 | 197 | 0.00 | 45 | 0.00 | 215 | 0.00 | 30 | 0.00 | 68 | 0.00 |
| snRNA | 2,245 | 0.09 | 12,537 | 0.08 | 1,862 | 0.15 | 8,234 | 0.08 | 920 | 0.06 | 2,527 | 0.03 | 448 | 0.04 | 1,134 | 0.01 |
| snoRNA | 104 | 0.00 | 134 | 0.00 | 166 | 0.01 | 253 | 0.00 | 87 | 0.01 | 120 | 0.00 | 989 | 0.09 | 1,428 | 0.02 |
| tRNA | 11,969 | 0.46 | 247,535 | 1.52 | 19,779 | 1.58 | 183,847 | 1.87 | 16,681 | 1.07 | 174,613 | 1.90 | 5,462 | 0.47 | 49,742 | 0.54 |
| unann | 2,495,922 | 96.55 | 13,691,523 | 84.19 | 1,136,866 | 90.75 | 7,556,930 | 76.83 | 1,479,737 | 94.96 | 7,453,740 | 81.00 | 1,102,285 | 95.41 | 6,445,195 | 70.46 |

### Sequence alignment and phylogenetic analysis

The mirBase (http://www.mirbase.org/) accession numbers for miR-1 and family members are shown in Table 4. Multiple sequence alignments were analyzed using Clustalx (1.81) software. A phylogenetic tree was constructed with the sequences obtained in this study and sequences of miR-1 precursor sequences from different species available in the miRBase data library using neighbor-joining in MEGA 7 software [39].

### Cell culture and luciferase assay

The 293T cell line used in this study was maintained in Dulbecco’s modified Eagle’s medium (DMEM) (Gibco, Waltham, USA) supplemented with 10% fetal bovine serum (Gibco), penicillin and streptomycin in an incubator with 5% CO_2_ at 37°C. Predicted binding sites were cloned and inserted into the pmirGLO vector (Promega, Madison, USA). For reporter assays, 150 ng of pmirGLO reporter vector and 50 nM miR-1 mimic (RiboBio, Guangzhou, China) were cotransfected into 293T cells using Lipofectamine 2000. No-mimic-treatment cells were used as a blank control, and cells carrying the pmirGLO-Hsp vector alone were used as the negative control. Firefly and Renilla luciferase activities were measured 48 h posttransfection by Dual-Luciferase Reporter Assay System (Promega). First, 100 μL of luciferase assay reagent II was added to each well, firefly luciferase activities were measured, and 100 μL Stop&Glo reagent was then added. Renilla luciferase activities were then measured. Firefly luciferase in the pmirGLO vector was used for normalization of Renilla luciferase expression. Treatments were assessed in triplicate, and transfections were repeated three times. Firefly luciferase activities were divided by Renilla luciferase activities for each experiment, providing the ratio.

### Synthesis and application of antagomir

Antagomirs, microRNA-specific antisense oligonucleotides. were synthetized by Dharmacon (http://dharmacon.gelifesciences.com). The microRNA-1 antagomir (Ant1) is the reverse complement of mature microRNA-1, and chemical modification was performed as described in a previous study [40] (5’-mC.*.mG.*.mC.mG.mC.mG.mC.mU.mA.mC.mU.mU.mC.mA.mG.mG.mU.mA.mC.mC.*.m U.*.mG.*.mA.*-Chl-3’). The “missense” (MsAnt) sequence (5’-mC.*.mG.*.mC.mU.mU.mU.mC.mG.mU.mG.mG.mU.mU.mC.mU.mG.mG.mU.mA.mC.*.m C.*.mU.*.mU. *-Chl-3’) was used as the negative control for the antagomir. [“*” is a phosphate backbone modification that was introduced to increase nuclease resistance and facilitate cellular uptake and bioavailability in vivo. “m” is a 2’-O-methyl (2’-OMe) modification, which reduces off-targeting. “Chl” is cholesterol, which can enhance gene silencing in vivo]. Noninjected ticks were used as a blank control group. To assess the specificity of the antagomir, we measured the levels of other microRNAs, such as microRNA-10. Antagomirs were microinjected into unfed adult female *Ha. anatolicum* at a dose of 400 μM in 0.5 μL. Every group consisted of 30 female ticks, and the groups were given a blood meal on a host at 24 h after microinjection.

### Statistical analysis

All data were analyzed with GraphPad 5 using Student’s t-test. Probability values of less than 0.05 were considered significant, and results are shown as the mean ± SEM.

## Results

### Small RNA Library Construction and Solexa Sequencing

To identify microRNAs involved in different stages of *Ha. anatolicum* development, four small RNA libraries pooled from eggs, larvae, nymphs and adults were constructed and sequenced using an Illumina HiSeq2000 high-throughput sequencer. We compared datasets from the four libraries with the repository of mature animal microRNAs to known microRNAs in miRBase21 (http://www.mirbase.org/). As a result, a total of 16,262,023, 9,835,555, 9,202,455 and 9,147,849 raw reads were obtained for the egg, larval, nymph and adult libraries, respectively. After removing low-quality reads, adaptors, and insufficient tags, 2,585,169, 1,252,678, 1,558,217 and 1,155,283 clean reads of 18-30 nt were obtained. Length distribution analysis showed that most reads were 21–30 nt long, with the highest percentage of reads (23.14%) being 28 nt long; 15.15% were 27 nt (Figure 1).

**Figure 1:**
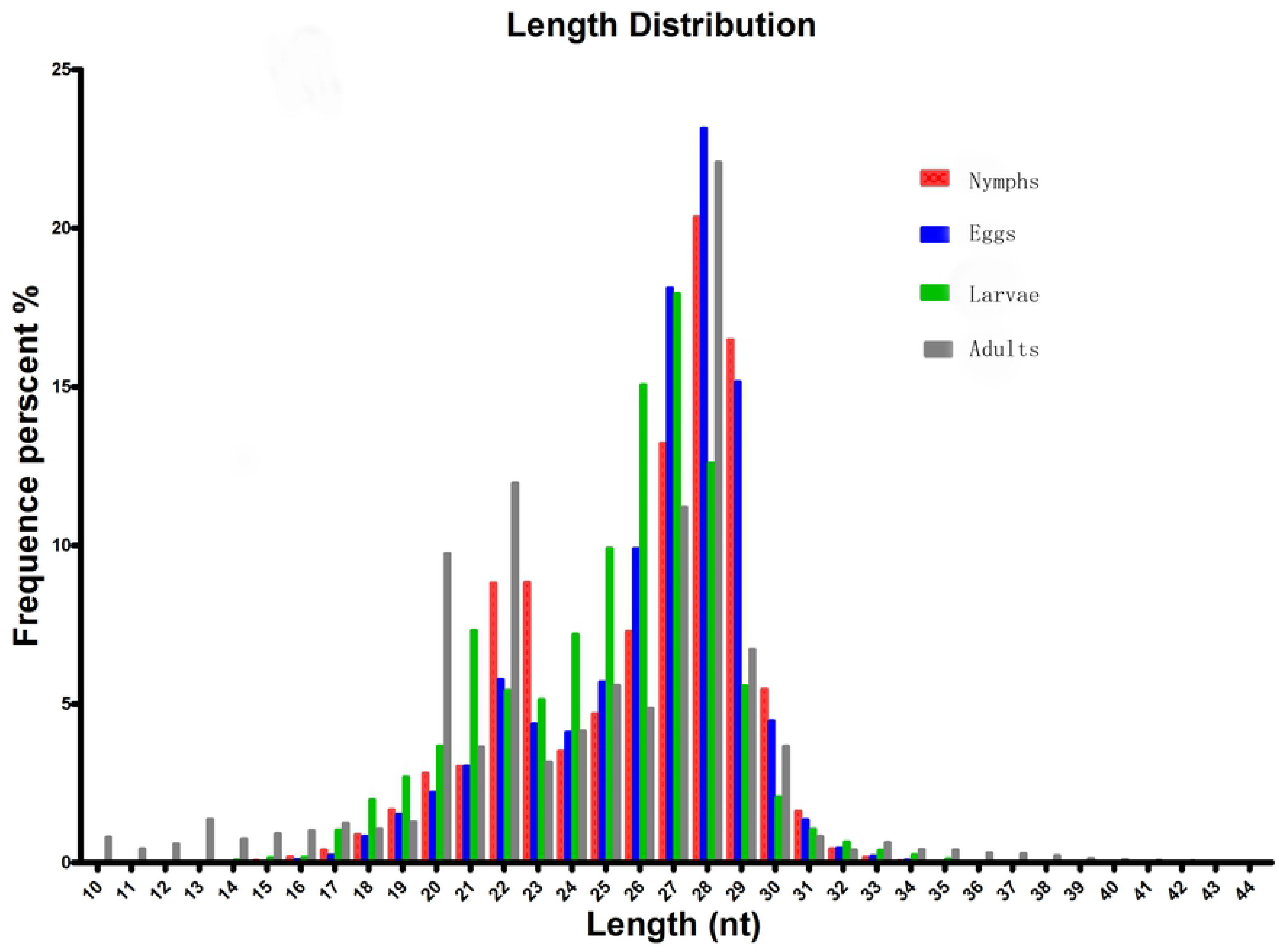
Length distribution and abundance of sequences in different developmental stages of *Hyalomma anatolicum anatolicum*. Contaminant reads from the fastq file were removed, and clean reads. Then, the length distribution of these clean reads was summarized. Normally, the length of small RNA is between 18 nt and 30 nt. The length distribution analysis is helpful to assess the compositions of small RNA samples. For example, miRNA is normally 21 nt or 22 nt, siRNA is 24 nt, and piRNA is 30 nt.

To assess the efficiency of high-throughput sequencing for sRNA detection, all sequence reads were annotated and classified through alignment with GenBank and Rfam databases. Of these sequences, which account for 1.29%, 2.86%, 1.72% and 1.92% of unique small RNAs (sRNAs) for egg larvae, nymphs, and adults, respectively, were perfectly mapped to the *Ixodes scapularis* genome (Accession: PRJNA34667) (Table 1). In total, 72.61% are common sequences to larvae and eggs, 48.89% between larvae and adults, 46.86% between eggs and adults, and 47.00%, 79.42% and 71.64% between nymphs and adults, nymphs and larvae, and nymphs and eggs, respectively; 9.67% miRNAs were common to all larvae, nymphs, and adults (Figure 2).

**Figure 2:**
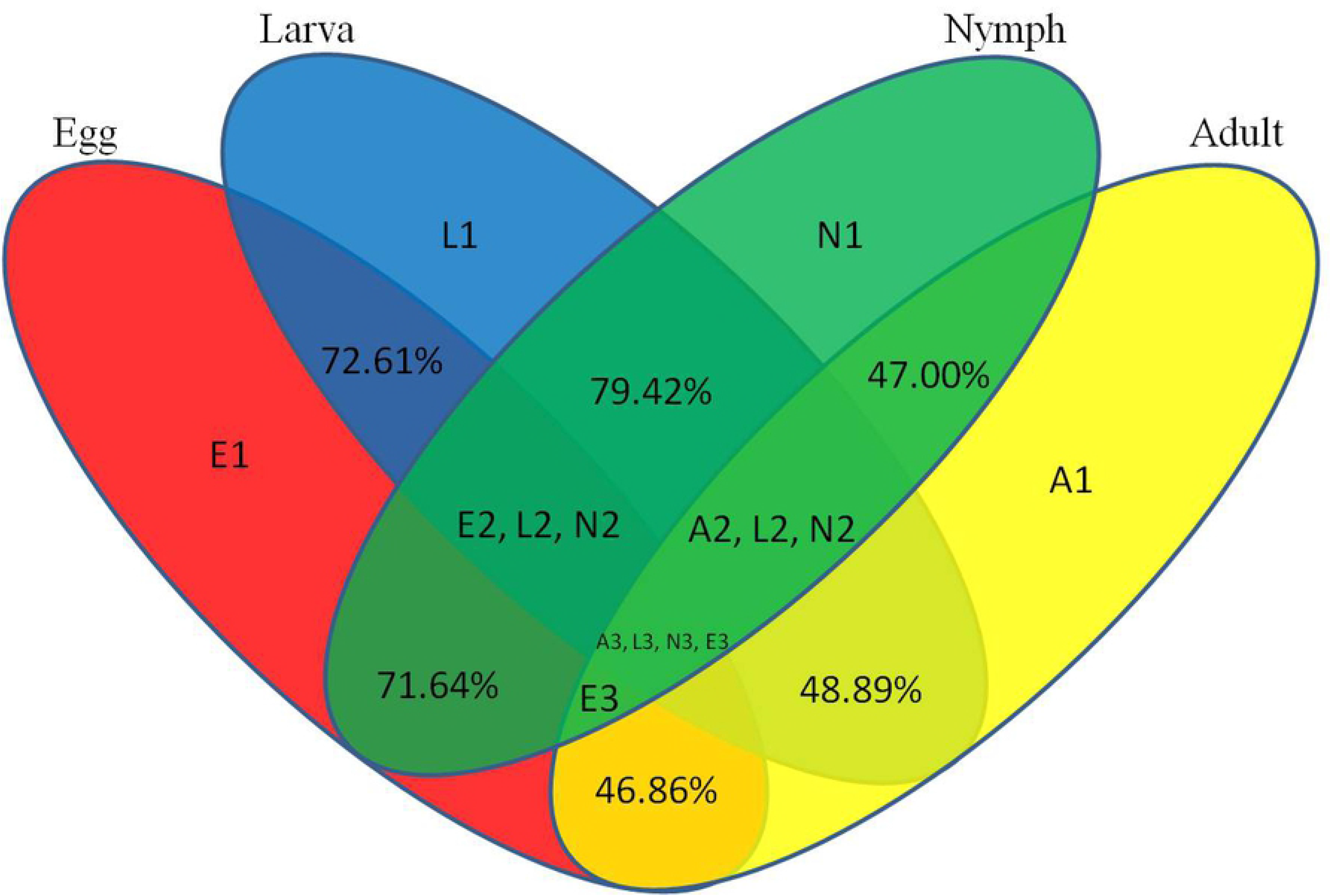
Common and specific unique tag sequences between samples. The percentages indicate common microRNAs between developmental stages. E1, L1, N1, and A1 indicate specific microRNAs in eggs, larvae, nymphs and adults, respectively. “E2 L2 N2” indicates common microRNAs in eggs, larvae and nymphs, respectively. “A2 L2 N2” indicates common microRNAs in adults, larvae and nymphs, respectively. “E3” indicates common microRNAs in adults, eggs, nymphs and adults. “A3 L3 N3 E3” indicates common microRNAs in all developmental stages.

### Known microRNAs and differential expression analysis

In this study, known microRNAs from *Ha. anatolicum* ticks were analyzed by miRBase21. The results showed a total of 232 known microRNAs in the egg stage, 1,051 in larvae, 1,122 in nymphs and 743 in adults. In eggs, miR-4175-3p and miR-4419b were predominately expressed, with more than 100,000 reads, and some microRNAs constituted 17.78% (260,875/1,467,411) of the total sequencing reads, suggesting that they are abundantly expressed during this period. The sequencing frequencies of 916 microRNAs were much lower than the 10 reads in larvae, but miR-184, miR-1 and miR-184b were predominately expressed. This was also observed at the nymph tick stage. However, in adults, miR-1-3p, miR-1, let-7-5p, miR-4486, and miR-84a were the most abundant, each with more than 100,000 reads. A total of 1,368 microRNAs displayed the lowest sequencing frequencies, with no more than 10 reads in larvae, nymphs and adults. In the four libraries, miR-1-3p, miR-1, and miR-4175-3p were detected with high abundance (Additional file 1). Compared with microRNA expression in various developmental stages, 987 microRNAs were significantly differentially expressed, with a p-value<0.01. When larvae were used as controls, 193 microRNAs were significantly differentially expressed in adults and 88 in eggs. Similarly, in nymphs, 355 microRNAs were differentially expressed. When using eggs as a control, 55 significantly differentially expressed microRNAs were detected in adult ticks and 74 in nymphs. If adults were used as a control, there were 222 significantly differentially expressed microRNAs in nymphs (Figure 3; Additional file 2). These microRNAs were mainly expressed at low levels in different developmental stages, such as miR-12-5p, miR-1357, miR-1193-5p, bantam-b, and miR-252b.

**Figure 3:**
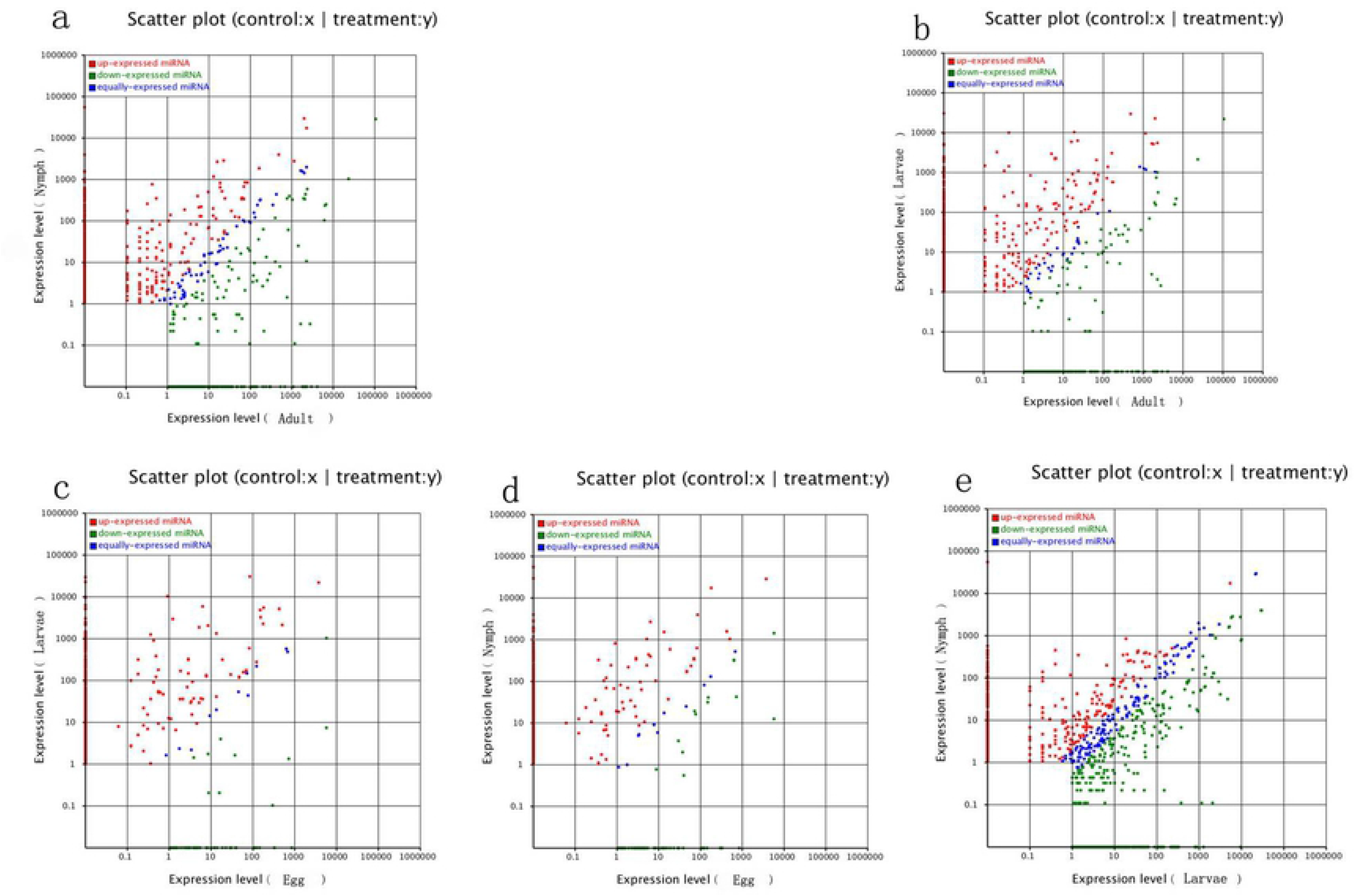
Comparison of differential expression levels of known microRNAs in different developmental stages. Each point in the figure represents a microRNA. The X-axis and Y-axis show the expression levels of microRNAs in the two samples. The red points represent microRNAs with ratios >2; the blue points represent microRNAs with 1/2<ratios ≤2; the green points represent microRNAs with ratios ≤1/2. Ratios = Normalized expression in the treatment/Normalized expression in the control. Panel a represents different expression between nymphs and adults; b represents different expression between larvae and adults; c represents different expression between larvae and eggs; d represents different expression between nymphs and eggs; e represents different expression between nymphs and larvae.

Currently, there are 49 known microRNAs in *Ix. scapularis*, a species belonging to the prostriate hard-tick lineage [41,42], but there are no known microRNAs identified for *Ha. anatolicum* or other metastriate hard-tick species. Our results indicated no known microRNAs in the egg stage but 46 in the larval stage, 45 in the nymph stage and 41 in the adult stage (Table 2).

**Table 2.**
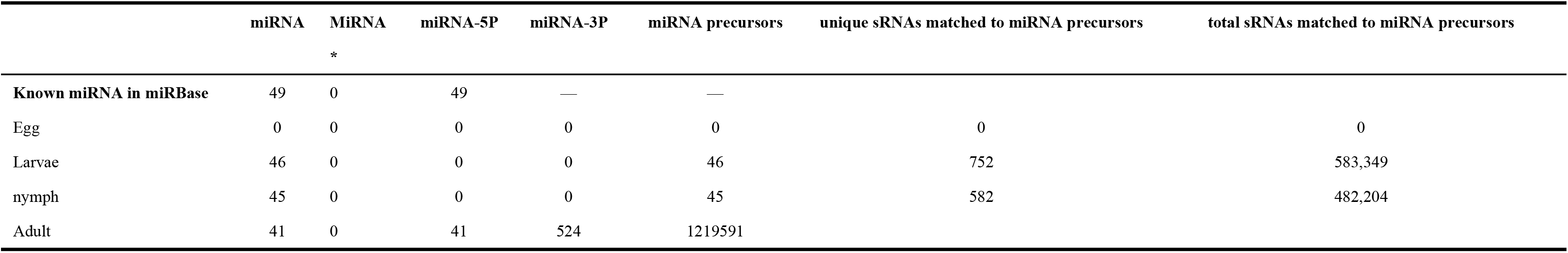
Known miRNA alignment and expression profile. Align sRNA to the miRNA precursor of corresponding species (if there is no precursor information of that species in miRBase, mature miRNA is ok) to obtain the miRNA count; If there is miRNA information of the species in miRBase, align sRNA tags to the miRNA precursor/mature miRNA of corresponding species in miRBase; if not, align sRNA tags to the miRNA precursor/mature miRNA of all plants/animals in miRBase, see Addition file 1.

### Identification of novel microRNA candidates

In addition to profiling known microRNAs, high-throughput sequencing has the advantage of revealing functionally important novel microRNAs that might not be detected using traditional methods. The unannotated unique sRNAs and total sRNAs are shown in Table 1. To determine whether these small RNA sequences are genuine tick microRNAs 17, 30, 16 and 10 potential novel microRNAs in eggs, larvae, nymphs and adults were examined. The number of total novel microRNA candidates was 1012, 4143, 2721 and 2130 in different developmental stages (Table 3). The characteristic hairpin structure of the microRNA precursor can be used to predict novel microRNAs. We used the prediction software Mireap to predict novel microRNAs by exploring the secondary structure, Dicer cleavage site and minimum free energy of unannotated small RNA tags that could be mapped to the genome. Mireap can be accessed from the following link: http://sourceforge.net/projects/mireap/ [34]. The lengths ranged from 20 to 24 nt, with a typical stem-loop structure and free energy ranging from −43.5 kcal mol^−1^ to −19.4 kcal mol^−1^ (Additional file 3).

**Table 3.** Map the clean tags back to genome by SOAP2[1] to analyze the expression and distribution of sRNA tags across the genome. If the genome sequence of the species is not available, the corresponding EST or one specified species with available genome sequence as the reference will also work. The characteristic hairpin structure of miRNA precursor can be used to predict novel miRNA. We developed Mireap (http://sourceforge.net/projects/mireap/), a prediction software to predict novel miRNA by exploring the secondary structure, the Dicer cleavage site and the minimum free energy of the unannotated sRNA tags which could be mapped to genome. Note: [1] Li R., Yu C., Li Y., et al. SOAP2.An improved ultrafast tool for short read alignment. Bioinformatics, 25 (15):1966- 1967.

| Type | Unique sRNAs | Mapping to genome | Total sRNAs | Mapping to genome | Number of unique novel miRNA candidates | Number of total novel miRNA candidates |
| --- | --- | --- | --- | --- | --- | --- |
| Eggs | 2,585,169 | 79,076 (3.06%) | 16,262,023 | 3,099,347 (19.06%) | 17 | 1,012 |
| Larvae | 1,252,678 | 29,211 (2.33%) | 9,835,555 | 1,036,333 (10.54%) | 30 | 4,143 |
| Nymphs | 1,558,217 | 20,176 (1.29%) | 9,202,455 | 533,540 (5.80%) | 16 | 2,721 |
| Adults | 1,155,283 | 15,034 (1.30%) | 9,147,849 | 2,099,735 (22.95%) | 10 | 2,130 |

### MicroRNA target gene prediction and GO enrichment

We screened microRNA 1 (miR-1), which exhibited a high expression level, as an important regulatory factor in ticks. To further understand the physiological functions and biological processes involving miR-1 during various developmental stages, target gene prediction was performed based on microRNA/mRNA interactions to provide molecular insight into processes. The MireapV0.2 software results revealed target genes involved tracheal system development, transcription factors, transmembrane receptors, and notum morphogenesis, among others (Additional file 4). GO, an international standardized classification system for gene annotations that provides insight into the molecular functions of genes in various biological processes [37], enrichment analysis was performed (http://geneontology.org/). In the cellular components category, there were 66 genes with a P-value≤1. Moreover, 57.50% of the genes clustered into the term organelles. Regarding molecular function, 76 genes were assigned; most were related to catalytic activity, with 51 (occupied 67.80%) annotated genes. Analysis of biological processes showed that 266 genes are involved in macromolecule metabolic processes or nitrogen compound metabolic processes, at 36.80% and 23.20%, respectively. Figure 4 illustrates the global analysis of GO enrichment of targets.

**Figure 4:**
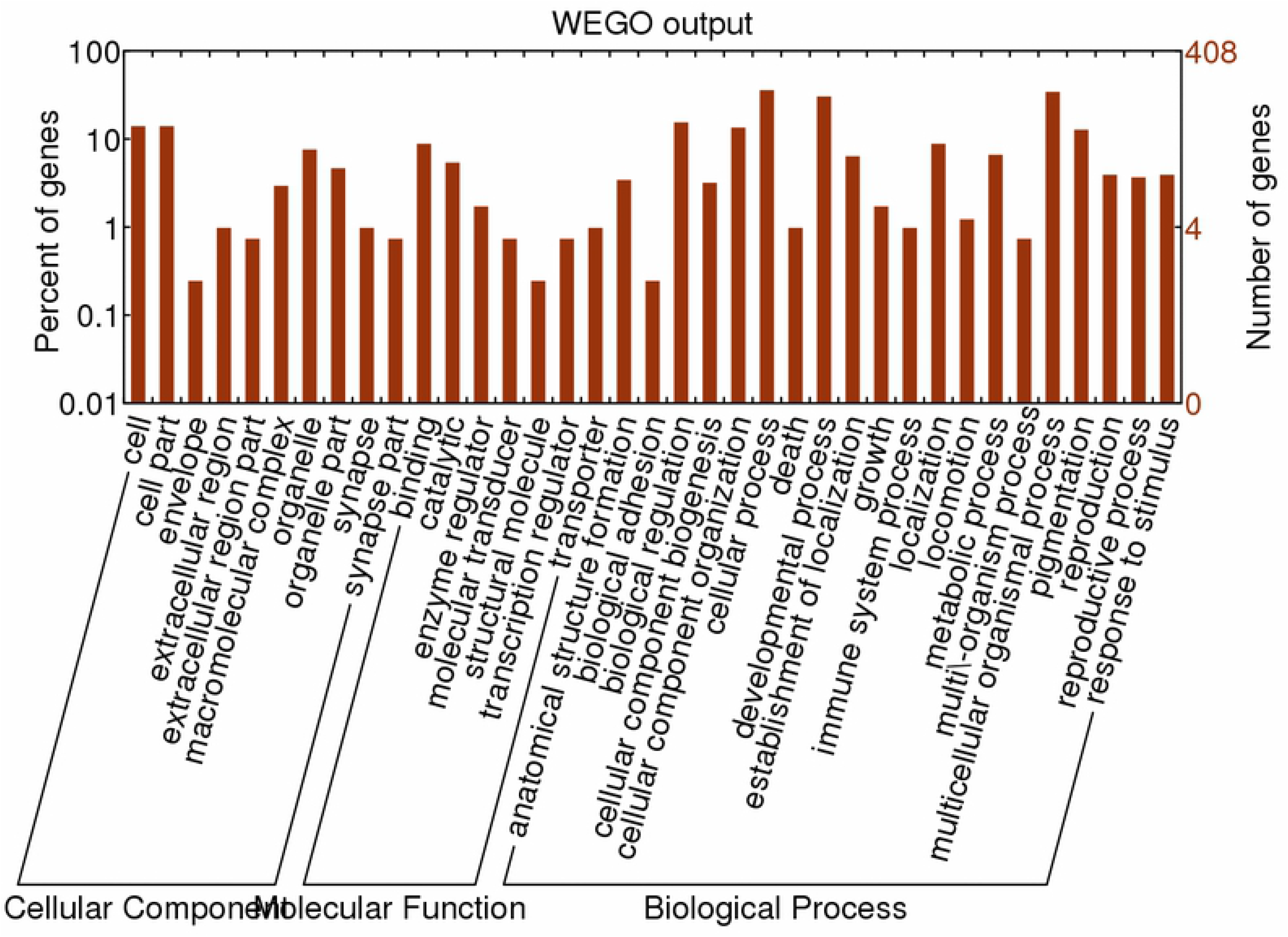
Partial GO classification annotated by Gene2GO for predicted target genes. The figure shows partial GO enrichment for the predicted target genes in terms of cellular components, molecular functions, and biological processes.

### Analysis of miR-1 conservation

In addition, we examined miR-1 hairpin precursor sequences in different species. The 3’-arm of the hairpin is highly conserved, though the many changes in the 5’-arm are fully consistent with the precursor hairpin structure. Other miR-1 sequences are short and very similar, and their genomic contexts can improve our ability to annotate and explore their evolutionary origins (Figure 5A). The genomic organization of miR-1 family members across phyla suggests that miR-1 is an ancestral microRNA. The characteristic hairpin structure of microRNA precursors can be used to predict microRNAs. Mireap [5] (http://sourceforge.net/projects/mireap/) was used to predict microRNA by exploring the secondary structure, Dicer cleavage site and minimum free energy of the unannotated sRNA tags mapped to the genome (Figure 5B).

**Figure 5:**
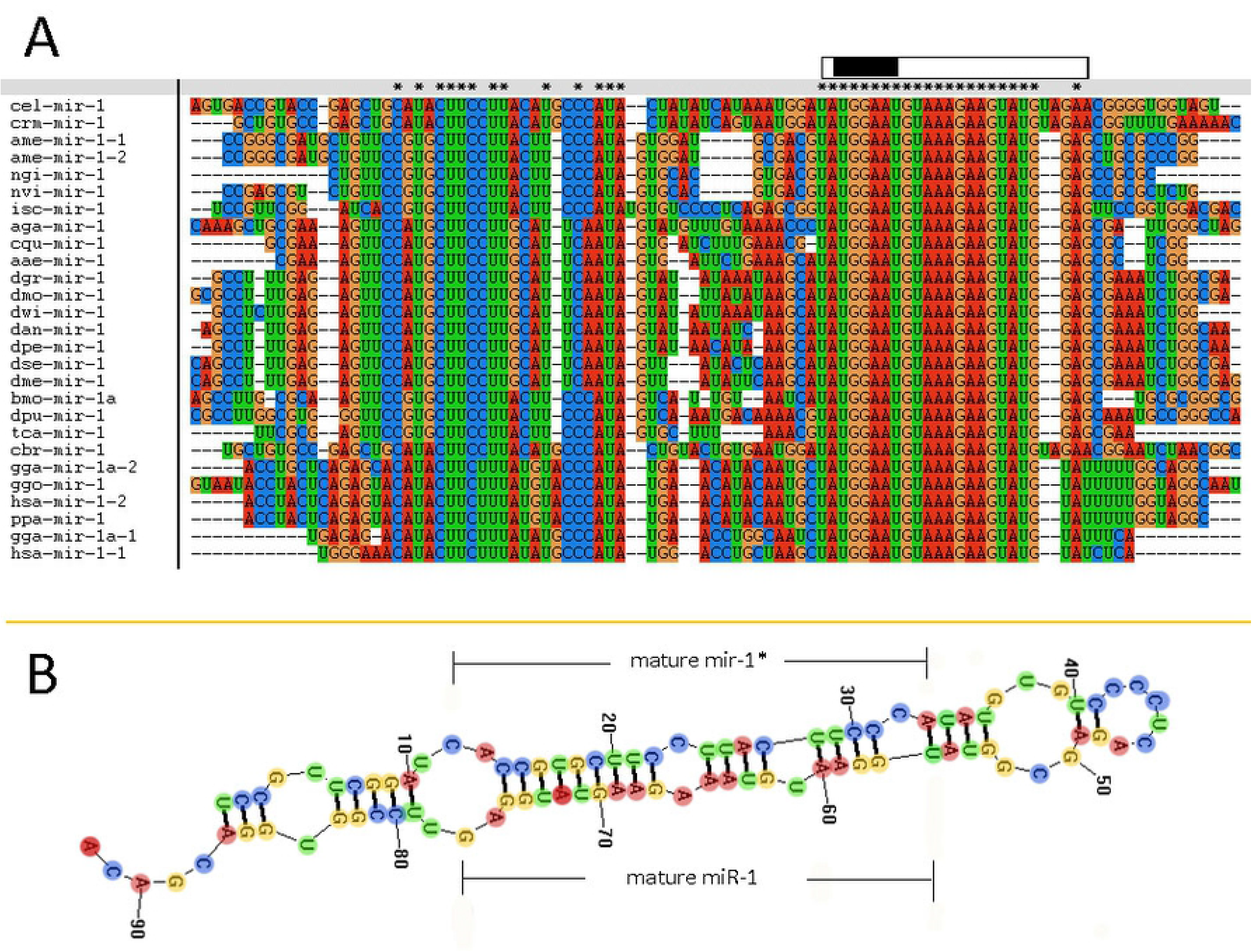
Comparative results of the nucleotide sequence of each species miR-1 (Clustal w software). “*”indicates conserved sequences. “partial precursor sequence” is the precursor sequence of miR-1. “Mature” denotes the mature sequence of miR-1. “seed region” is the seed region of miR-1.

We further investigated the evolutionary pattern of this gene family in vertebrates and found that it has been generated by multiple replications of the ancestor gene, including two duplications as a whole and one fragmental replication, with mutation and deletion of certain genes in some species (Figure 5 and Figure 6). Analysis of the phylogenetic distribution of the miR-1 family in various species indicated that it is an ancient gene family originating in the urochordate *Caenorhabditis elegans.* In contrast, *Drosophila melanogaster* contains only a single copy of the miR-1 gene, though the genomes of vertebrates contain more than one copy.

**Figure 6:**
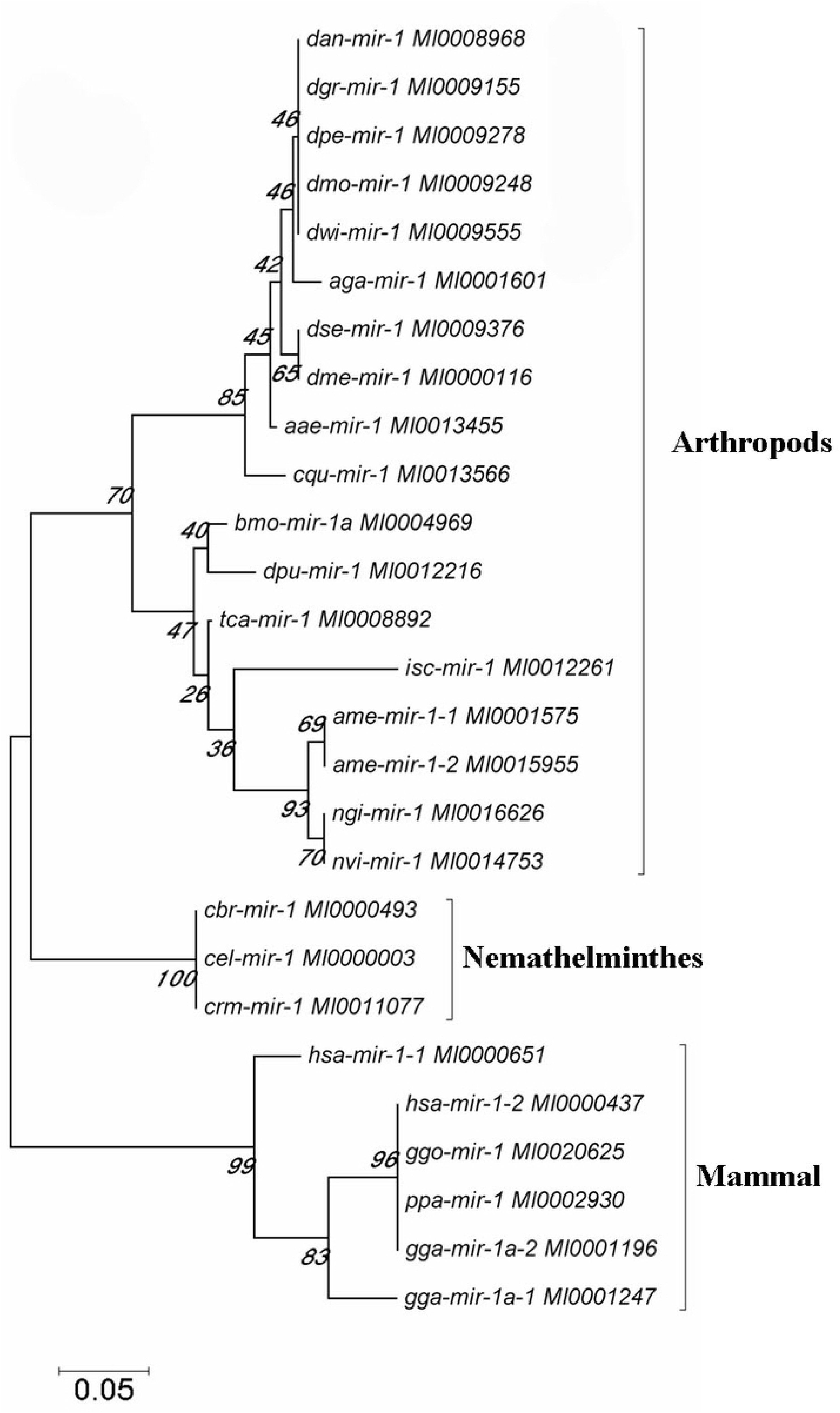
The phylogenetic tree was constructed for the miR-1 gene in selected species by neighbor-joining (MEGA 4.0 software) with maximum parsimony and 500 bootstrap replicates. Reference nucleotide sequences were selected by BLAST searches of the NCBI nt database.

### miR-1 targets Hsp60

The putative target gene (Hsp60) of miR-1 was identified using RNAhybrid, and the complete sequence of Hsp60 was used to predict binding sites. Computational analysis revealed many potential binding sites in Hsp60, but only one predicted result was within the 7-mer seed sequence sites (we chose the definition of the canonical seed binding site; the seed sequence is generally defined as 2-8 bases) in Hsp60 (Table 4). Based on these results, one binding site was cloned and inserted downstream of Renilla luciferase in the pmirGLO vector, which was then cotransfected into 293T cells with miR-1 mimics. According to luciferase reporter assays, only one site (position 342) resulted in 48.50% luciferase activity compared with that of the negative control and no-mimic control (Figure 7); the other site resulted in no significant difference compared with that of the control. Therefore, Hsp60 is a potential target of miR-1 in vitro. Expression analysis of miR-1 and Hsp60 for various developmental stages and tissues in ticks was then performed.

**Table 4:**
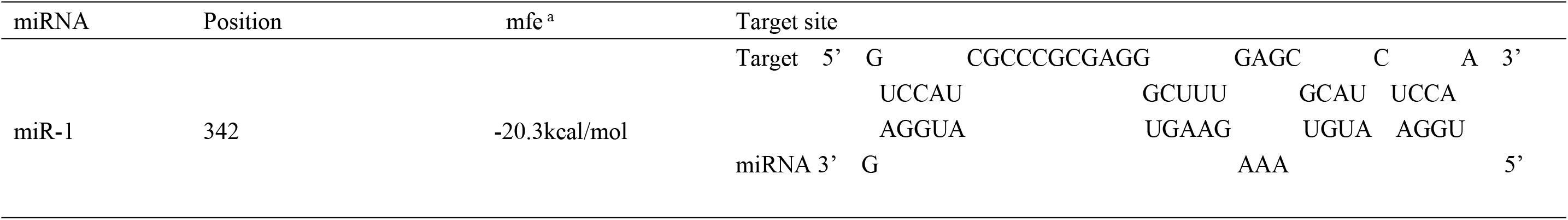
Putative miR-1binding sites of HSP60 ^a^ minimum free energy (mfe) values based on RNAhybrid 2.2 prediction

**Figure 7:**
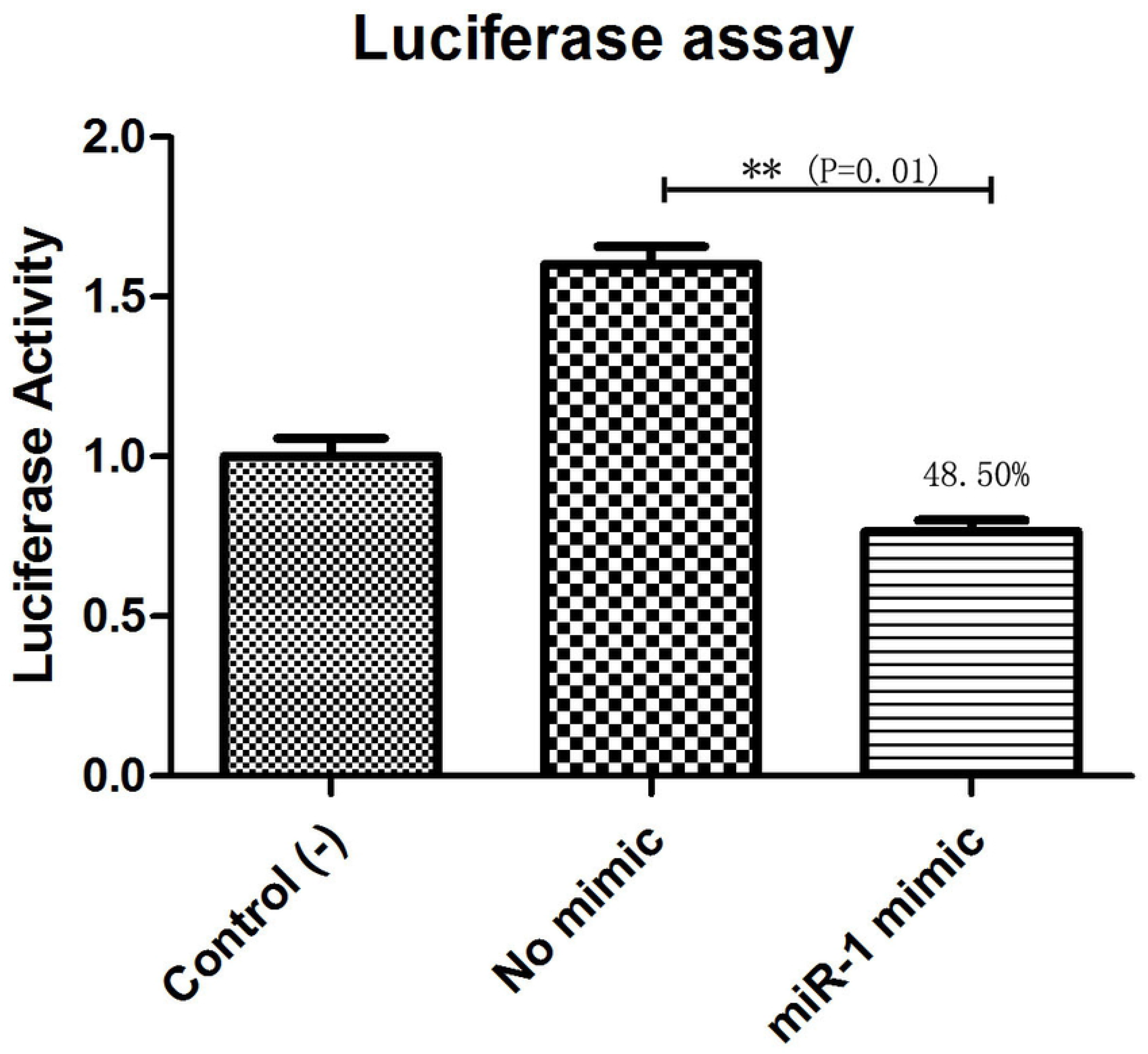
Hsp60 is a target of miR-1. Dual luciferase reporter assay results are represented as the mean±SEM of triplicate samples. The recombinant plasmid pmirGLO containing the Hsp60 gene was transfected into 293T cells. Expression of Hsp60 was measured by qPCR. Data are presented as the mean±SEM of triplicate samples. **P≦0.01 (Student’s t-test)

To further confirm Hsp60 as an authentic miR-1 target gene in vivo, we conducted phenotype rescue experiments using Hsp60 RNAi in female ticks with an Ant-1 background; we speculated that RNAi-mediated knockdown of the physiologically relevant target of miR-1 would alleviate the adverse phenotypes caused by miR-1 depletion. Indeed, coinjection of Ant-1/dsRNA partially alleviated these phenotypes. Furthermore, the Ant-1/dsRNA female tick body weight significantly increased after a blood meal compared with that of dsHSP60 ticks. Thus, Hsp60 is an authentic target of miR-1 in vivo.

### Expression analysis of miR-1 and Hsp60 for various developmental stages and tissues in ticks

To investigate the tissue- and developmental stage-specific expression of miR-1 in ticks, we measured expression levels of mature miR-1 at different developmental stages (egg, unfed larvae, fed larvae, unfed nymphs, fed nymphs, unfed adults and fed adults) and in various tissues (the midgut, ovary, and salivary glands) of unfed and fed female ticks using real-time PCR. Expression analysis of mature miR-1 in different developmental stages showed that miR-1 expression peaked at the partially fed female stage (Figure 8a). In addition, expression of Hsp60 was highest at the engorged adult stage but was drastically reduced in eggs (Figure 8a). Analysis of different tissues indicated higher mature miR-1 levels in salivary glands than in other studied tissues in unfed female ticks; in fed female ticks, levels were highest in the epidermis (Figure 8b). To further investigate the potential function of miR-1 in adult female ticks, we silenced miR-1 by Ant-mir1: each unfed female tick was microinjected with Ant-mir1 or MsAnt, and after 48 hours, real-time PCR was performed to assess the silencing efficiency of Ant-mir1. The results showed that miR-1 expression levels decreased to 64.67% after injection of Ant-mir1 compared with MsAnt and noninjection controls (Figure 9) (t=5.800, df=4). In this experiment, the host blood (rabbit) and male ticks were also examined by qPCR after injection of Ant-mir1, with insignificant levels in the rabbit blood. However, the expression level was higher in males than in females, and the expression level of miR-1 was significantly different in engorged females injected with water and Ant-mir1 (Figure 10).

**Figure 8:**
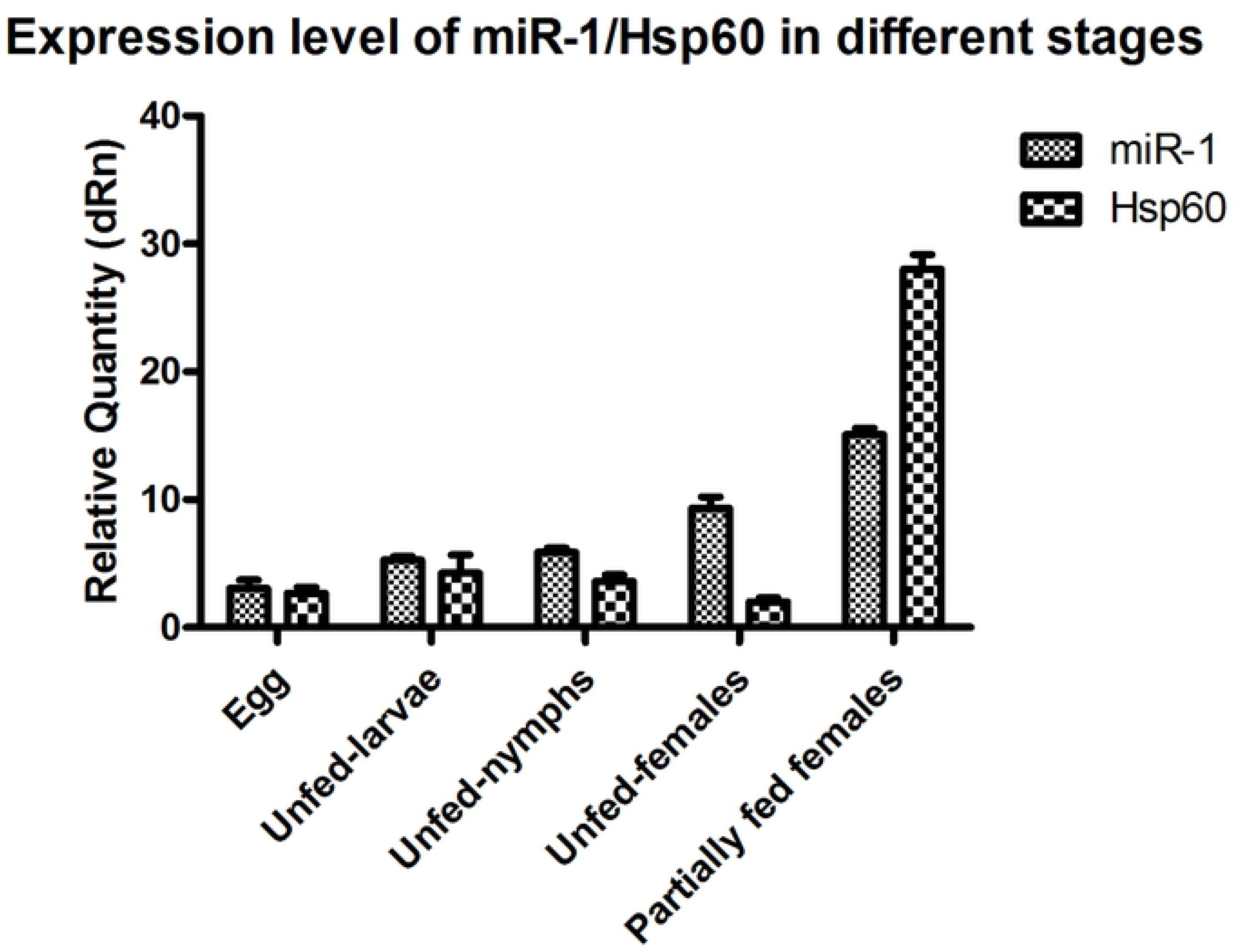
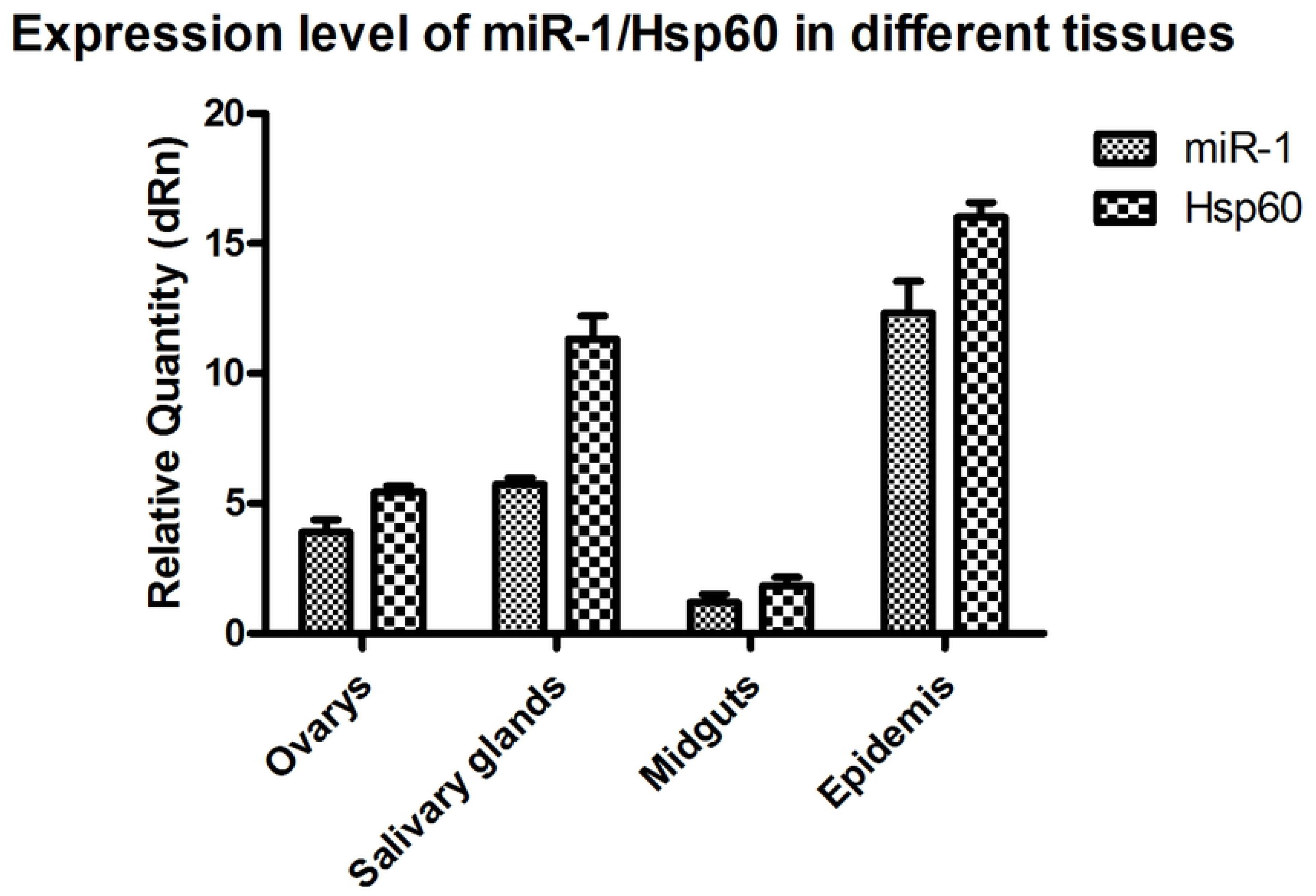
miR-1 and Hsp60 expression in different developmental stages and tissues. a Relative expression of Hsp60 and miR-1 was analyzed in eggs, unfed larvae, unfed nymphs, unfed adult females and partially fed adult females. b Relative expression of Hsp60 and miR-1 in the ovary, salivary glands, midgut and epidermis of fed adult female ticks. Data represent three biological replicates with three technical replicates and are shown as the mean ± SEM.

**Figure 9:**
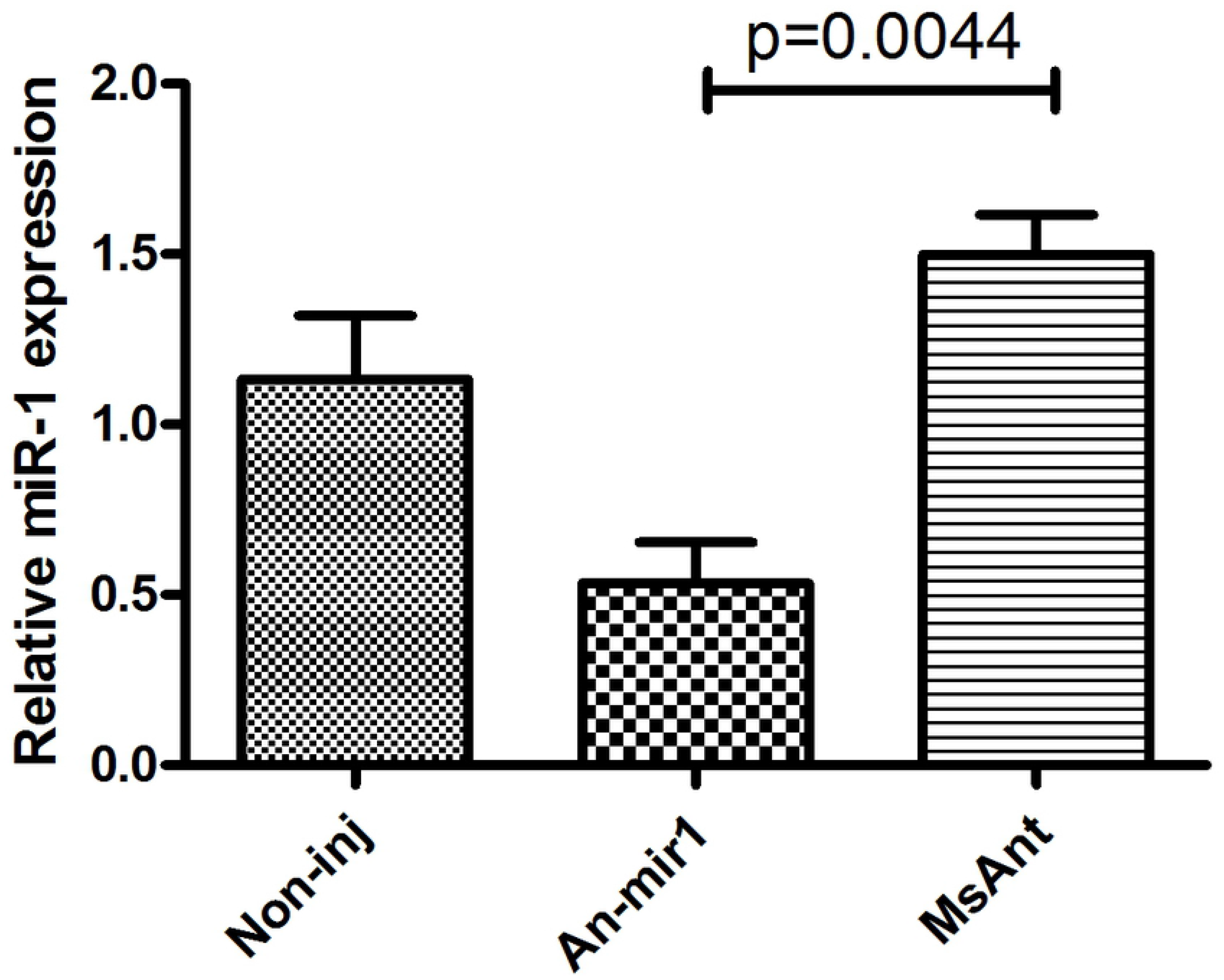
Relative mature miR-1 expression in fed female ticks treated with the miR-1 antagomir. Data represent three biological replicates with three technical replicates and are shown as the mean ± SEM.

**Figure 10:**
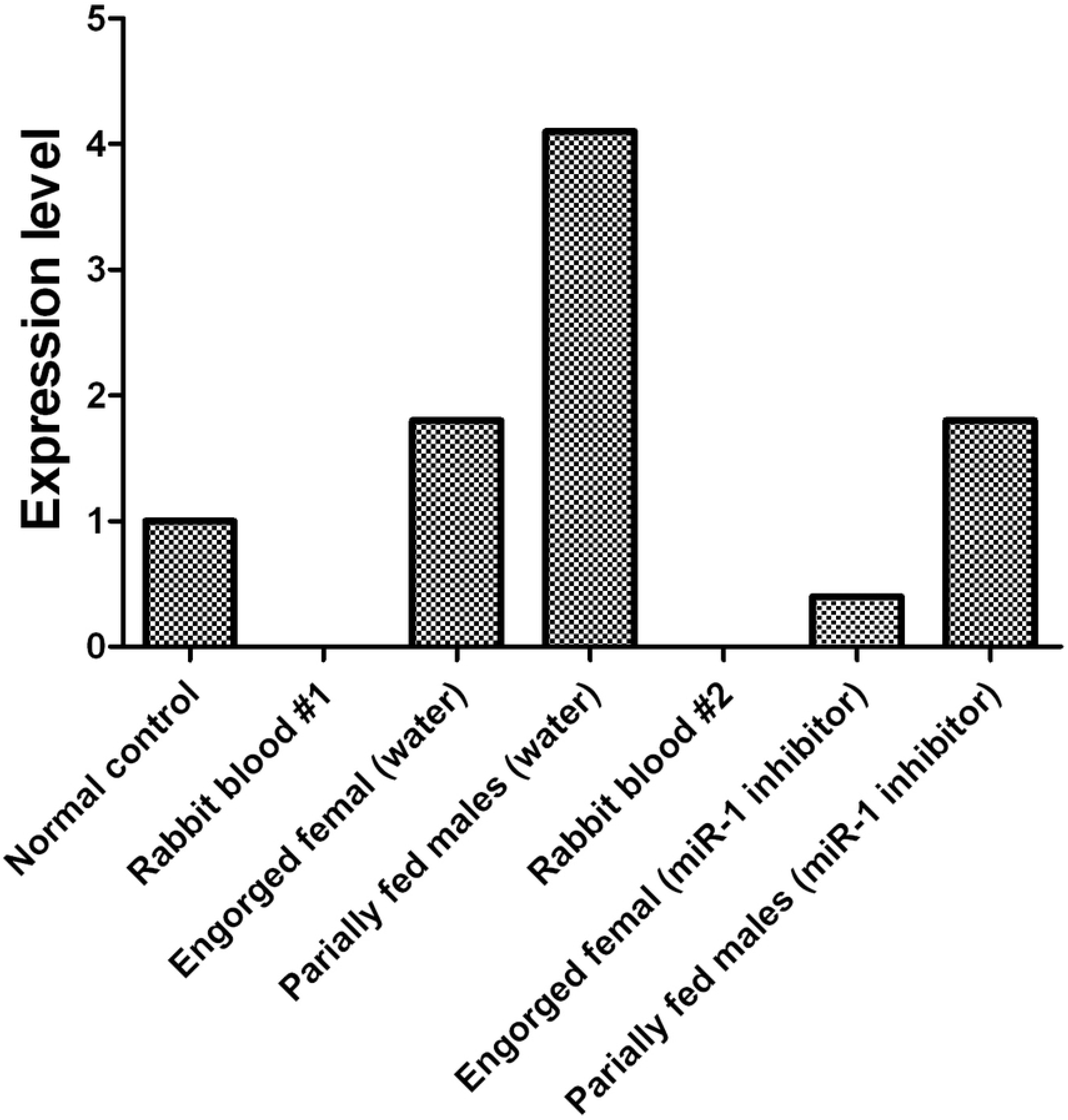
Before and after inhibition of miR-1, dynamic expression during tick engorgement was determined. Rabbit blood and water were used as controls.

### Physiological effects of microRNA inhibition on ticks

Phenotypic manifestations were also monitored in female ticks after treatment with Ant-1, with a significant reduction in blood-feeding time after Ant-1 treatment compared with the MsAnt control group observed. According to analysis of the change in feeding, compared to control groups, fed female ticks injected with Ant-1 showed a much faster increase in the duration of feeding for an average of eleven days. An average of twelve days after engorgement was found for the control group, and most female ticks began to lay eggs and lost weight. The weight of Ant-1-treated females was 81.00±1.9 mg, but the MsAnt- and noninjection groups weighed 127.20±2.1 mg and 119.48±2.0 mg, respectively, after engorgement, with average egg weights of 52.15±3.2 and 48.96±3.1, respectively. However, the spawning and mortality rates were not obviously affected (Table 5).

**Table 5:**
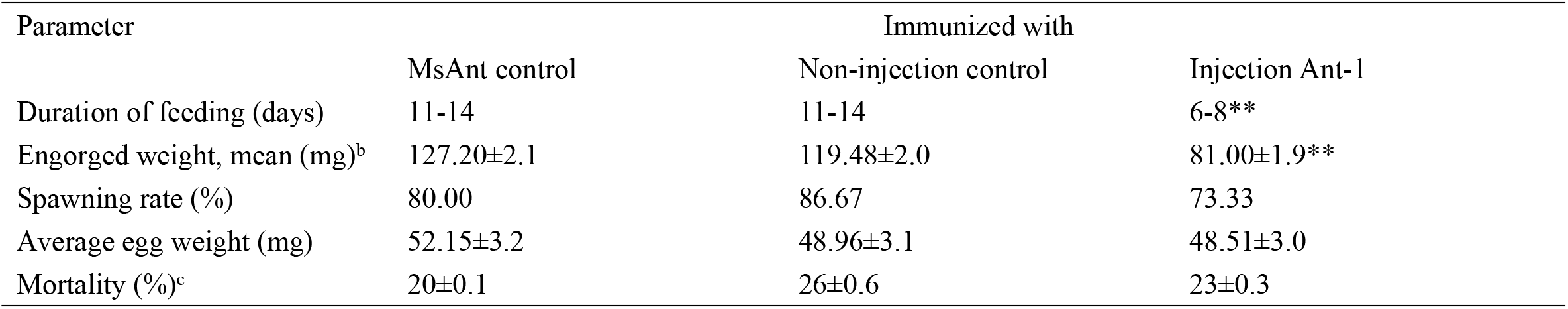
Effect of vaccination with miR-1 inhibitor the ticks feeding ^a^ Statically significant (P<0.05) as calculated by Students *t* test. a Values are expressed as follow: average ± standard deviation. b Dead ticks were excluded, Calculated as batch average (batch weight/number of adult ticks). c Mortality rate was calculated as the number of deaths during and after feeding period

### MicroRNA analysis of the Hsp60 gene

After inhibition of miR-1, no significant changes in physiological indicators, including engorgement time, were detected. It may be that other members of the microRNA family compensate for the loss of miR-1. To confirm this finding, ticks with miR-1 inhibition were used as the model strain with Hsp60 as the target gene for microRNA family prediction. The results showed that other microRNAs acting on the Hsp60 gene include mir-5, mir-994, and mir-969, among others. Therefore, RT-PCR was employed to detect expression levels of other microRNAs when miR-1 was inhibited; upregulation was found (Figure 12), with miR-5, which recognizes the same site as miR-1, being most significantly upregulated.

**Figure 11:**
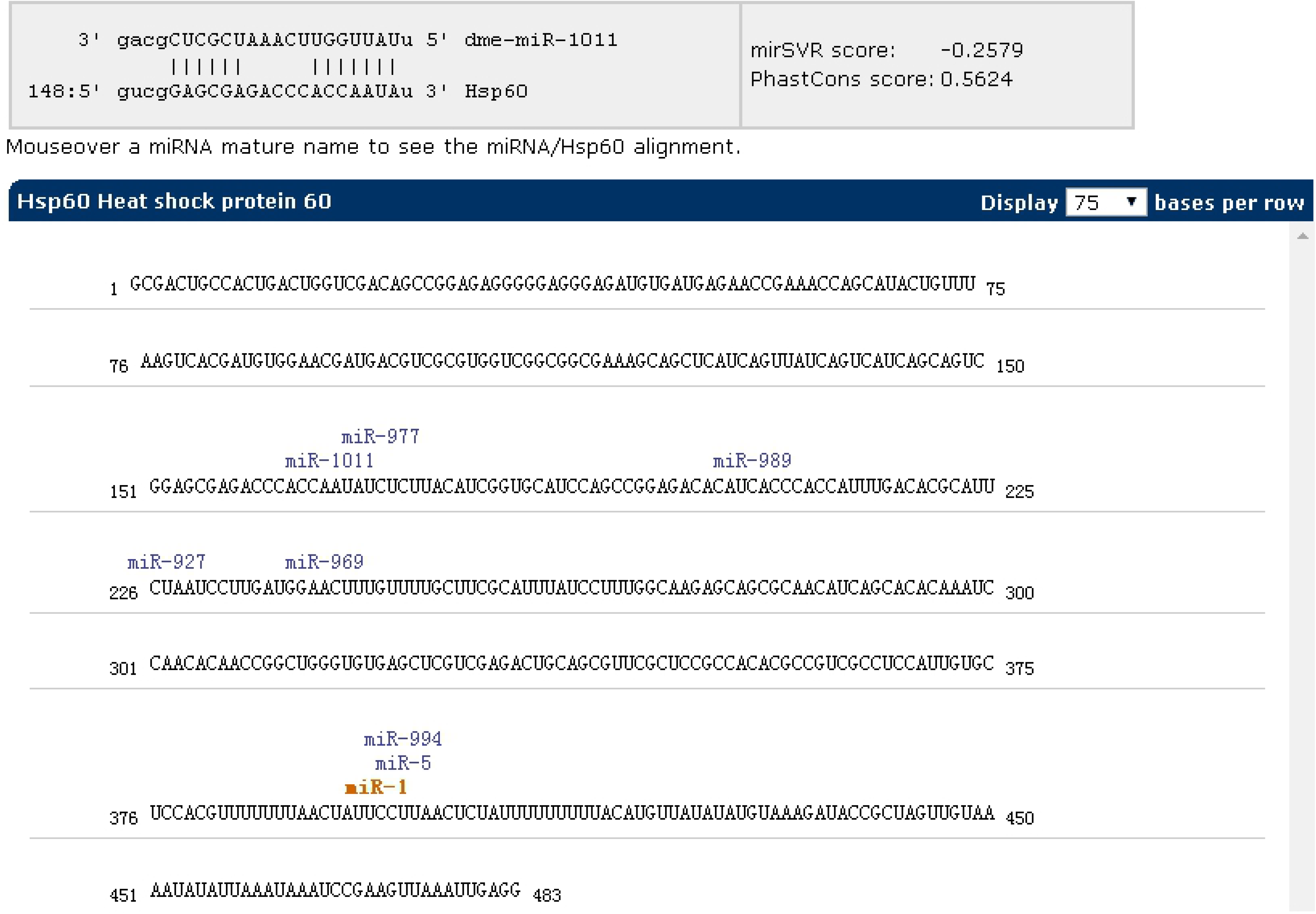
miR-1 inhibition as the model strain and Hsp60 as the target gene for microRNA family prediction using miRWalk (version 3.0).

**Figure 12:**
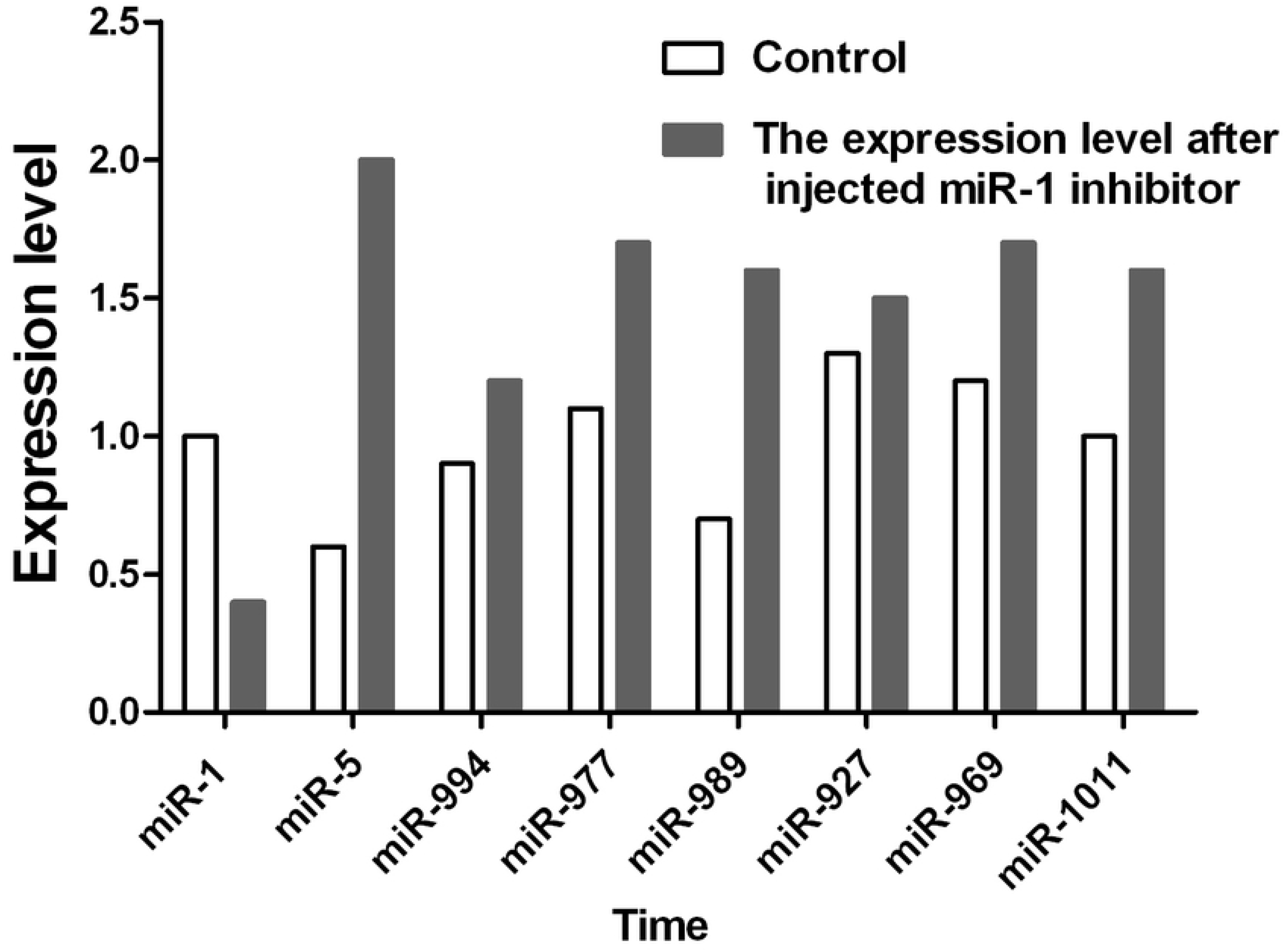
The expression level of the miR-1 family was detected by RT-PCR when miR-1 was inhibited.

## Discussion

Small RNAs are a key molecules in organisms that induce gene silencing and have important roles in the regulation of cell growth, gene transcription and translation. Small RNA digitalization analysis based on HiSeq high-throughput sequencing uses SBS sequencing, which can decrease loss of nucleotides caused by secondary structure. This approach is also beneficial because of the small sample quantity requirement, high throughput, and high accuracy via a simply operated automatic platform. Such analysis can result in millions of small RNA sequence tags in one run, comprehensively identify small RNAs of certain species under certain conditions, predict novel miRNAs and assemble small RNA differential expression profiles between samples; overall, it is a powerful tool in small RNA functional research.

We analyzed small RNAs at different developmental stages in *Ha. anatolicum* by HiSeq sequencing. Assessment of fragment length to detect the peak of small RNA length distribution from ticks can help to determine relevant types of small RNA, such as miRNA of 28 to 22 nt, siRNA of 24 nt and piRNA of 30 nt. In this study, 89% of the segments were in the range of 19 ~ 24 nt (Figure 1), conforming to high-throughput sequencing quality control [30] and providing a foundation for the reliability of later relevant data analysis. The distribution of small molecular RNAs of different lengths during tick development may help to uncover important aspects of tick physiological functions [43]. For example, miRNAs were the most abundant small RNA, with 1.29% unique to eggs, 2.86% to larvae, and 1.92% unique to adults. It is approximately 95% for the unannotated small RNA in different developmental stages (Table 1), indicating a very complex regulatory system. As shown in Table 1, miRNAs comprised a relatively large proportion of all the small RNAs annotated in larval ticks, indicating that miRNAs are involved in the regulation of gene expression in the early stage of tick development. This result can provide a basis for screening functional miRNAs by analyzing specific and common sequences as well as a target for the study of specific regulatory molecules during tick development. Common miRNAs function in maintaining the physiological characteristics of ticks at different developmental stages (Figure 2). Nevertheless, specific expressed miRNAs may have important species-specific functions and/or host and pathogen specificity. These results provide a reliable basis for targeted study of miRNAs, and the differential miRNAs identified are of great significance for investigation of function.

Moreover, known miRNAs expressed in different samples were statistically analyzed to determine differences in expression level between two samples, and the differentially expressed miRNAs commonly expressed in the two samples were compared using a scatter plot (Log2 -ratio) (Figure 3). Analysis of miRNA expression in different developmental stages showed approximately 800 miRNAs to be differentially expressed in these samples. For example, we detected 854 differentially expressed miRNAs between larvae and adults, 894 between nymphs and adults, 754 between eggs and nymphs, and 694 between eggs and larvae. In addition, using eggs as a control, Bantam and Let-7 miRNA and miR-133 showed large differences in various developmental stages. However, differential expression of miRNAs between adult and nymph and larval ticks was mainly found in the Let-7 family, such as Let-7 g, Let-7i, miR-144 and miR-1495. Based on these results, different types of miRNA are expressed among different stages of development in the same tick species. It is significant that miR-1 is present not only at different developmental stages of ticks but also in abundance (millions of transcripts). This indicates that miR-1 plays an important physiological function in maintaining normal physiological metabolism in ticks. Previous reports have confirmed that miR-1 is involved in many biological processes in animals. In mammals, miR-1 regulates the development of myocardial cells, and abnormal regulation leads to heart diseases such as myocardial infarction, arrhythmia, and cardiac hypertrophy [26]. Furthermore, the functions of miR-1 were assessed by GO analysis, revealing that miR-1 plays an important role in molecular binding and transcriptional regulation and is mainly involved in biological regulation, development, immune response, reproductive process and cell death (Figure 4). Dysregulation of miR-1 may inhibit apoptosis or cause cell proliferation. Although multisequence alignment indicated significant differences in the nucleotide precursor sequences of miR-1 from different species, the mature sequence is conserved with regard to primary structure. Specifically, the “seed region” is identical among different species, though the “B region” and “C region” exhibit a difference of 2-3 bases (Figure 5). This is because the miRNA binds to target genes via the seed region, allowing specific binding between the miRNA and target gene; variable regions of mature sequences emphasize the diversity of these sequences. Such conservation in miRNAs has been applied for the early diagnosis of cancer, species identification, and pathogen detection as well as other purposes. The results also suggest both specificity and diversity in target mRNAs. In addition, we found showed significant differences in nucleic acid sequences for precursors of miR-1 and related species. However, these species clustered into three main branches in phylogenetic trees (Figure 6), demonstrating large differences among arthropods, Nemathelminthes and primates. For example, ticks, mosquitoes and Drosophila display high identity in the arthropod group. In particular, only the “C” region was identical, with 1-2 base differences in mature sequences of miR-1 from different species; the “seed region” was also identical. Therefore, miR-1 is an ancient regulatory molecule that plays an important role in the evolution of species.

In this study, it was shown that miR-1 is expressed at various developmental stages in tick development and in different tissues, and miR-1 expression was upregulated with the growth and development of ticks. In particular, the expression level was significantly higher during engorgement than starvation (Figure 7A), and miR-1 was mainly expressed in the epidermis and midgut (Figure 7b). These results indicate that miR-1 not only maintains normal physiological function but also has a leading role in tick muscle development. It is possible that with the engorging process, the tick’s muscles gradually extend to accommodate the filling of the midgut. To confirm this, miR-1 was inhibited via injection of an inhibitor, and the change in physiological indices was significant (Table 5). As time is required for inhibitors to function in living animals, the expression level of miR-1 in ticks was tracked to determine the optimal feeding time after injection, and miR-1 expression was downregulated significantly after 16 h (Figure 8). Ticks were released on the surface of animals for a long time and then collected, and total RNA was extracted to detect miR-1 expression (Figure 9).

Previous studies have shown that heat shock proteins (Hsps) are directly related to the immune protection caused by the invasion of foreign agents in a host [44]. A previous report showed that a 63-kD symbiotic protein was abundant in probacterial symbionts in specialized cells of aphid hemolymph called mycetes; the protein is highly homologous (88.7% similar) to GroEL, a member of the Hsp60 family in *Escherichia coli* [45] and plays a role in viral spread. Our GO functional enrichment of microRNA analysis indicated that Hsp60 is a key target gene of miR-1 (as shown in Table 4 and Figure 10). miR-1 regulates the transcription of Hsp60, participating in the formation of specific tissues and organs during development. In other studies, the Hsp60 gene was associated with the development of the hematopoietic system and hematopoietic stem or progenitor cell proliferation, differentiation, maturation, and tumorigenesis [46–51]. Therefore, it will be helpful to explore the functions of miR-1 to understand the biological characteristics of Hsp60 and its expression (Figure 7).

miR-1 is a highly conserved microRNA molecule, and to avoid the influence of rabbit-derived miR-1 on our results, miR-1 was detected in rabbit blood. As no miR-1 was found in rabbit blood as a control, rabbit-derived microRNA would not affect tick physiology. In fact, expression was significantly lower than that in the control group in the miR-1 inhibition analysis and the inhibitory effect was significant. Thus, in ticks, miR-1 might regulate Hsp60 to resist damage by foreign agents. The findings also indicate that the binding of animal miRNA to target mRNA depends not only on the seed region but that the “B” and “C” regions determine the specificity for the interaction in different species.

Previous studies have confirmed that miRNAs can be specifically expressed in high abundance in animal reproductive tissues, such as miR-449a, miR-465c, miR-202, and miR-547 [52]. In addition, miR-34, miR-469, miR-465, and miR-101 are differentially expressed during testicular development [53]. Therefore, miRNA plays an important role in spermatogenesis, fertilized egg development and gametic differentiation [54, 55]. In this study, the expression level of miR-1 in male ticks was significantly higher than that in female ticks, suggesting that miR-1 is key in the maturation of sperm. This conclusion needs to be confirmed by further experiments, which may provide ideas for studies on tick sexual reproduction and parthenogenesis. Our miRNA inhibition experiments confirmed that after mating with female ticks injected with miR-1 inhibitor, the spawning rate of males dropped to 40%, and the incubation rate was only 10%. Further observation of the developmental morphological characteristics of hatchling ticks revealed severe aberrations in secondary development after miR-1 inhibition. Furthermore, physiological indices such as engorgement weight and time and mortality of female ticks were analyzed, and it was found that the engorgement time of ticks injected with miR-1 inhibitor was significantly shorter than that of the control group (time= 7±1 d for the experimental group, time=12±2 d for the control group). Regardless, there was no significant difference in other indicators (as shown in Table 5). To analyze the reasons for the lack of difference in physiological changes, we predicted other miRNAs possibly regulating Hsp60 and found that miR-5, miR-994, miR-969, miR-927, miR-989, miR-977, miR-1011 and miR-1 are common families that regulate the function of Hsp60 (Figure 9). We speculate that when miR-1 expression is inhibited, these family members will compensate for the loss. To verify this, qPCR was applied to detect expression of other miRNA clusters after miR-1 inhibition, which showed that other miRNAs were significantly upregulated after miR-1 was significantly inhibited. In fact, the expression level of miR-5, which has the same binding site as miR-1, was 4 times the normal level. miR-989 was also significantly upregulated (Figure 10). These results indicate that miR-5 plays an equally important role as miR-989 in the regulation of Hsp60 by miR-1. When miR-1 was inhibited, the two miRNAs preferentially compensated for the loss of function.

In conclusion, miR-1, a small noncoding RNA present in lower to higher animals, is highly conserved that is expressed to varying degrees in different stages of tick development and between tissues, with relatively stable expression. miR-1 plays an important role in tick development. However, considering that multiple genes are regulated by the same factors, loss of miR-1 function is compensated for by other miRNAs from the same family. Nonetheless, this compensation is not complete. In the early stage of development, the damage caused by abnormal gene expression may be temporarily negligible, but this damage cannot be effectively compensated for in later stages of development. Therefore, aberrant expression of miR-1 in ticks seriously affects their fecundity, especially egg hatching. This also leads to deformities later in development.

## Acknowledgments

All listed authors made substantial, direct, and intellectual contributions to this work and approved its publication.

**Addition file 1:** Differential expression of known microRNAs between developmental stages. A is a pair of developmental stages in the differential expression analysis; B column is the microRNA name; C and D are adult total reads; E and F are true expression levels of microRNA; G and H (*-std) are normalized expression levels of microRNA in a developmental stage; I (fold-change (log2*/*)) is fold change of microRNAs in the pair of developmental stages, with negative numbers indicating downregulation and positive numbers upregulation; J is the P-value reflecting the significance of microRNA differential expression between developmental stages, whereby a smaller value indicates greater significance of the difference in microRNA expression between developmental stages and the last column (sig-label) “**”: fold_change (log2)>1 or fold_change (log2)<−1, and P-value<0.01. “*”: fold_change (log2)>1 or fold_change (log2)<−1, and 0.01 <=P-value<0.05. None: Others.

**Addition file 2:** When no microRNA information of the species was available in miRBase21, small RNA tags were aligned to the microRNA precursor/mature microRNA of all plants/animals. The sequence and count of microRNA families (no specific species) found in the samples.

A is the microRNA name (no specific species); B is the count of microRNAs of a family in the developmental stage; C is the sequence of a microRNA with the highest count in the family.

**Addition file 3:** The long sequence indicates microRNA precursor information in the order of sequence, name and long; The parentheses indicate microRNA precursor information in the order of hairpin structure, structure and MFE; “*** “and a short sequence indicate mature microRNA information in the order of sequence, name and length; “…” and a short sequence indicate information of matched sRNA tags in the order of sequence, ID, length, count.

